# Tangerine: A Python framework for dynamic gene regulation analysis from transcriptomic time series

**DOI:** 10.64898/2026.07.17.739167

**Authors:** Tanmayee Narendra, Gabriele Schweikert

## Abstract

**Motivation:** Time-series single-cell transcriptomics enables the study of dynamic gene regulation. However, standard computational tools frequently aggregate temporal data into static, dense topologies, obscuring the precise regulatory rewiring that drives developmental transitions. Further, navigating the inherent noise of statistical inference without losing biological interpretability remains an important bottleneck.

**Results:** We present Tangerine, a Python framework for the dynamic reconstruction and interactive exploration of time-varying gene regulatory networks. Tangerine integrates time-constrained metacell aggregation with regularized linear modelling and non-parametric correlation to infer dynamic topologies. To solve the interpretability gap, it features a browser-based visual analytics engine. Tangerine empowers researchers to track macroscopic gene module evolution, interactively filter effect sizes, and link topological rewiring directly to raw transcriptomic evidence.

**Availability and implementation:** Tangerine is implemented in Python and Plotly Dash. The code is available on Github at https://github.com/ntanmayee/tangerine.

## Introduction

Gene regulation is a highly dynamic process, and the precise rewiring of transcription factor (TF) networks drives cell fate and disease progression (Levine and Davidson (2005); Badia-i Mompel et al. (2023)). Within these broader networks, TFs and target genes frequently organize into tightly co-regulated clusters called gene modules, which activate or deactivate synchronously to drive specific biological programs. While recent computational advances have successfully transitioned single-cell analyses from descriptive atlases toward inferring dynamic, causal regulatory relationships (Bravo González-Blas et al. (2023); Wang et al. (2023); Kalfon et al. (2025)), extracting explicitly interpretable mechanisms remains a bottleneck. State-of-the-art inference tools increasingly capture these temporal dynamics; however, they frequently output mathematically dense networks, or encode regulatory logic within implicit hidden layers (Dimitrov et al. (2026)). As a consequence, researchers face an interpretability gap, where advanced statistical inference produces densely connected networks, which commonly obscure the specific, actionable biological switches that define developmental transitions. Further, computational inference of regulatory networks is inherently noisy, making it difficult to define rigid thresholds for true biological interactions (Chen and Mar (2018); Pratapa et al. (2020)). Researchers need tools that allow them to track continuous temporal effect sizes rather than relying on binary edge predictions. We argue that computational inference should not be an opaque endpoint, but rather the starting point for interactive biological exploration.

To address this interpretability gap, we present Tangerine, an interactive visual analytics framework designed explicitly for time-series single cell RNA-seq data. By integrating metacell aggregation with time-specific regularized linear modelling and non-parametric correlation, Tangerine reduces the memory footprint compared to unaggregated single-cell data, allowing execution on standard node-based workstations. This approach provides an extensible framework to infer dynamic, time-resolved topologies. Through a multi-scalar visual interface, users can navigate from macroscopic gene module evolution down to the raw transcriptomic distributions of a single shifting TF-gene interaction. Ultimately, Tangerine serves as an exploratory complement to statistical inference, empowering researchers to visually validate temporal regulatory rewiring without writing custom code.

## Methods

Tangerine is a Python framework comprising a computational backend for network inference and a browser-based visual analytics frontend. This architecture strictly decouples data processing from visualization (See Figure 1A for an overview).

**Figure 1.**
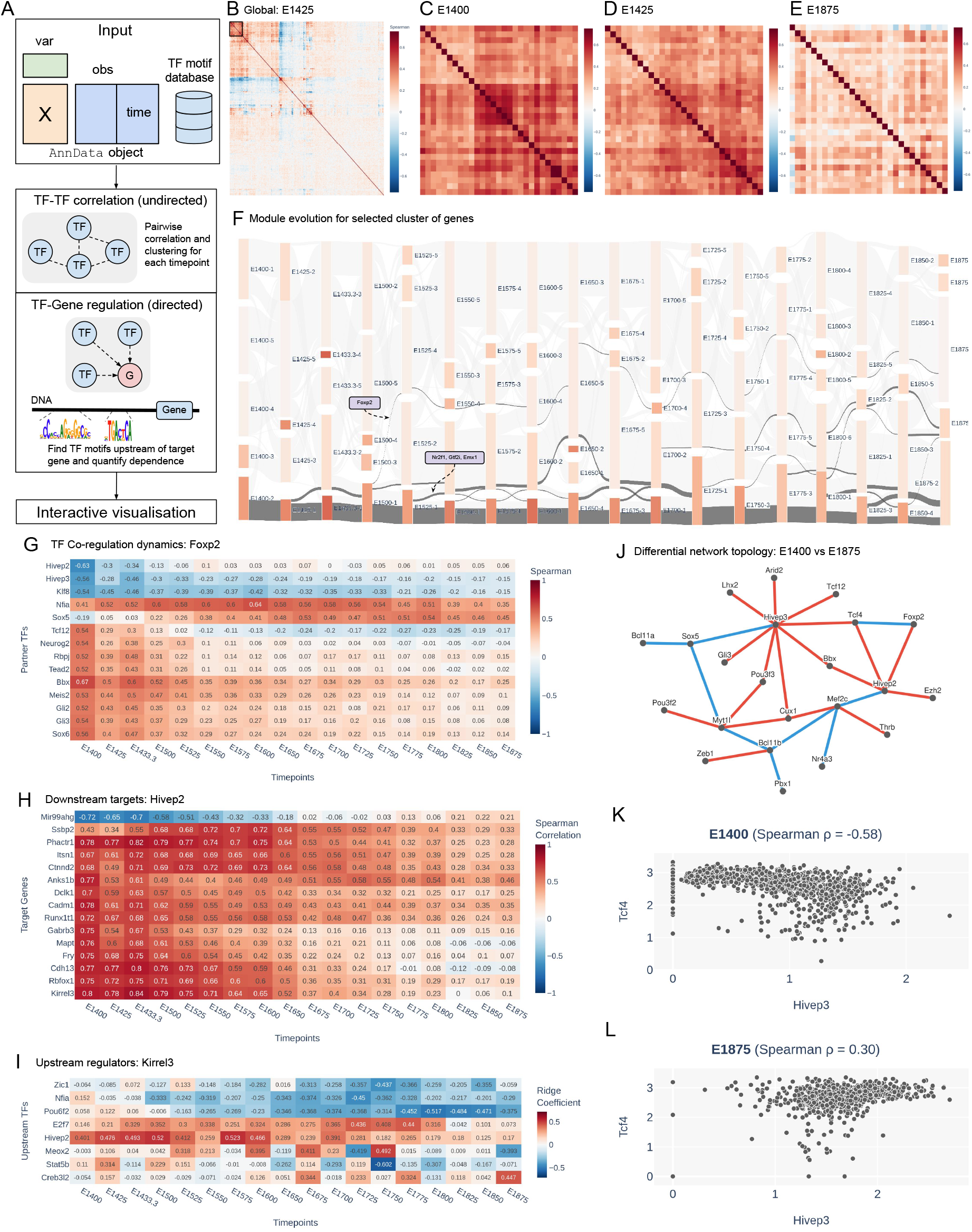
Tangerine provides a unified visual analytics framework for dynamic GRN inference. (A) Schematic of the computational pipeline, integrating sequence-based motif priors and time-resolved transcriptomic data to infer undirected (TF-TF) and directed (TF-Gene) regulatory topologies. (B) Global TF-TF Spearman correlation heatmap at timepoint E14.25, with a user-selected, tightly co-regulated sub-cluster highlighted. (C–E) Magnified views of the selected sub-cluster at E14.00, E14.25, and E18.75. Row and column orders are fixed to the clustering order in E14.25 to enable visual comparison of modules across time. (F) Interactive alluvial diagram tracking the selected sub-cluster and breakdown of the grouping across the time-course. (G–I) Targeted Dynamics views enabling granular inspection of regulation. (G) Spearman correlation profile of *Foxp2* with other co-regulatory TFs. (H) Directed regulatory influence of *Hivep2* on candidate downstream targets; all displayed target genes have the *Neurod2* binding motif upstream of their transcription start site (TSS). (I) Regulatory TFs driving the target gene *Kirrel3*. All TFs shown have their binding motif upstream of *Kirrel3*. (J–L) Differential Topology view capturing rewiring. (J) Differential network mapping highly changing TF interactions between timepoints E14.0 and E18.75, autonomously highlighting *Hivep3* as a rewiring hub. (K, L) Raw metacell expression scatterplots validating the inversion of the *Hivep3* -*Tcf4* edge

### Data Preprocessing

Single-cell transcriptomic data is inherently sparse; filtering genes based on dropout usually discards biologically relevant but lowly expressed genes. To deal with this zero-inflation, Tangerine aggregates cells into metacells independently within each timepoint, utilizing either the K-means or SEACells (Persad et al. (2023)) algorithms. This time-constrained aggregation prevents the artificial blending of different developmental states. Genes are then filtered based on their metacell dropout rates.

To establish a structural prior and reduce false-positive edges, the framework extracts DNA sequence data upstream of target gene transcription start sites. While this prior defaults to a 10 kb upstream window, this parameter is fully customizable, allowing users to incorporate distal enhancers or narrower promoter regions based on the specific genomic context of their organism. These regions are scanned for candidate TF binding motifs using GimmeMotifs (van Heeringen and Veenstra (2011)), restricting the set of potential regulators to those with physical binding capacity.

### Dynamic Inference

Tangerine applies two inference strategies independently to each timepoint. First, to model directed TF-gene causality, time-specific regularized linear models (Ridge, Lasso, or ElasticNet) are fitted to quantify the influence of candidate TFs on target genes. Regularization mitigates the multicollinearity inherent to co-expressed transcription factors (Hastie (2009); Kamimoto et al. (2023)).

Second, undirected TF-TF co-regulatory modules are inferred via non-parametric Spearman correlation. To track temporal rewiring, hierarchical clustering is applied to every timepoint specific correlation matrix. Next, optimal module assignments per timepoint are identified by using Ward’s minimum variance method (Ward Jr (1963)).

### Interactive Visual Analytics

To bridge the interpretability gap, Tangerine provides a browser-based visual analytics frontend that enables threshold-agnostic exploration of temporal networks. The interface allows users to navigate between macroscopic network topology and underlying raw transcriptomic data across three coordinated analytical views:

1. **Global Topology and Module Evolution:** A global TF correlation heatmap (Figure 1B) is linked to an alluvial diagram (Figure 1F). This allows users to visually select specific TF subsets in a given timepoint and trace their module membership across the time-course. Further, the selected TF subset can be compared in detail across user specified timepoints (Figure 1C-F). In these zoomed-in views, the ordering of genes is fixed to the global TF heatmap for all three timepoints.
2. **Targeted Dynamics:** Users can query specific target genes or regulator TFs to generate dynamic, clustered heatmaps of individual regulatory axes. Interactive sliders allow users to define correlation or regression coefficient thresholds to filter transient noise (Figures 1G-I).
3. **Differential Topology:** This component computes the change in Spearman correlation between any two TFs, at two user-selected timepoints. This allows the user to visually inspect TF-TF correlations with highest changes at the selected timepoints. The resulting topology is rendered dynamically using a constraint-based physics engine (Figure 1J). Selecting a TF-TF edge in the graph displays the underlying metacell expression distributions of the TFs in the selected two timepoints, linking graph topology directly to transcriptomic evidence (Figure 1K-L).

### Implementation Details

Tangerine is implemented in Python. The computational backend uses scanpy (Wolf et al. (2018)) for single-cell data handling, scikit-learn (Pedregosa et al. (2011)) for regularized regression, and networkx (Hagberg and Conway (2020)) for graph construction. To ensure low-latency data retrieval during interactive exploration, data is serialized into Parquet format, and inferred topologies are exported as GML files. The interactive frontend is built using the Plotly Dash framework, integrating the dash-cytoscape library to render the physics-driven network layouts in the browser.

## Results

To demonstrate Tangerine’s utility in resolving dynamic regulatory transitions, we applied the framework to a publicly available scRNA-seq dataset of mouse development (Qiu et al. (2024)), restricted to the intermediate neural progenitor major cell cluster. Data was downloaded from the CZ CELLxGENE Discover platform (Program et al. (2025)).

### Data Processing and Efficiency

The dataset was filtered to include only embryonic timepoints containing at least 10,000 cells. The final dataset comprised 282,095 cells across 18 timepoints (from embryonic day E14.0 to E18.75). Analysing dense time-series data of this scale typically imposes substantial memory constraints on standard hardware. Using K-means aggregation, the data was compressed into 519-1,340 representative metacells per timepoint. Tangerine completed the inference pipeline for this dataset in 2 hours and 32 minutes, with the first 47 minutes used for standard data pre-processing steps such as normalisation and computation of PCA and UMAP components. The peak memory usage was 68 GB.

### Macroscopic Module Evolution

Without prior biological seeding, Tangerine’s ‘Global Topology’ view successfully captured the macroscopic restructuring of the neurogenic regulatory network. At the E14.25 timepoint, we observe a tightly correlated cluster of 30 transcription factors (Figure 1B). This cluster includes *Neurog2*, expressed by progenitors poised to exit the cell cycle (Florio et al. (2012)); *Tcf12*, a known marker of neural progenitors (Uittenbogaard and Chiaramello (2002)); *Neurod1*, which directs fate decisions and maintains progenitor identity (Tutukova et al. (2021)); and *Foxp2*, a regulator of neural differentiation (Tsui et al. (2013)). This selected cluster of genes can be inspected in detail (Figure 1D) in comparison to two other timepoints (Figure 1C & E). Using the interactive Sankey plot to track this module through time, we observe that *Foxp2* exits this cluster at E15.0 (Highlighted in Figure 1F). Furthermore, a smaller sub-cluster of genes departs at E15.25; notably, this cluster includes *Nr2f1* and *Gtf2i*, which are primary drivers of the neurodevelopmental disorders Bosch-Boonstra-Schaaf Optic Atrophy Syndrome (BBSOAS) and Williams Syndrome, respectively (Bertacchi et al. (2020); Barak et al. (2019)). This demonstrates the framework’s ability to computationally isolate relevant temporal regulators from gene module dynamics.

### Resolving Granular Regulatory Rewiring

To investigate these transitions, the ‘Targeted Dynamics’ view was utilized to query specific TF rewiring across the time course. Figure 1G visualizes the correlation profile of *Foxp2*, isolating distinct positively and negatively co-regulated TF partners across the time-course. Additionally, Figure 1H displays the downstream target genes of *Hivep2*, revealing the progressive reduction of its regulatory influence across the time-series. Conversely, Figure 1I shows the regression coefficients of upstream TF regulators of target gene *Kirrel3*, where *Hivep2* has a prominent positive coefficient in the beginning of the time course. *Kirrel3* is known to be involved in neuronal migration, axonal fasciculation, and synapse formation (Hisaoka et al. (2018)).

The ‘Differential Topology’ view was then used to inspect the most highly divergent TF rewiring events between E14.0 and E18.75. The differential network revealed *Hivep3* as a highly volatile hub gene (Figure 1J). Querying *Hivep3* revealed a significant inversion in its regulatory relationship with *Tcf4*, a known driver of early neurogenesis (Flora et al. (2007)). Selecting the *Hivep3* -*Tcf4* edge projected the underlying metacell distributions, validating the topological shift with transcriptomic evidence. At E14.0, during peak neurogenesis, *Hivep3* and *Tcf4* exhibited a strong negative correlation (Spearman *ρ* = *−*0.58), consistent with the strict mutual exclusivity characteristic of early progenitor maintenance (Figure 1K). By E18.75, this antagonism degrades, resulting in a mathematical inversion of the edge weight (Spearman *ρ* = 0.30) (Figure 1L). This transition from an opaque network edge to a quantifiable degradation of mutual exclusivity demonstrates Tangerine’s capacity to identify and mathematically validate developmental regulatory switches.

## Conclusion

In the rapidly evolving landscape of single-cell transcriptomics, extracting actionable regulatory mechanisms from time-series data remains a persistent bottleneck. Tangerine addresses this challenge by re-framing gene regulatory network inference not as a static computational endpoint, but as an interactive, visually driven analytical process. By strictly decoupling computationally intensive operations such as sequence-based motif scanning, regularized regression, and hierarchical clustering, from the frontend visualisation interface, Tangerine provides a highly responsive environment for threshold-agnostic exploration.

A core philosophy of the framework is that statistical inference in biological systems is inherently noisy. Rather than enforcing rigid mathematical thresholds that may discard subtle biological signals, Tangerine empowers researchers to integrate their domain knowledge directly into the analytical loop. Through the resolution-adaptive visual engine, users can fluidly manipulate effect-size thresholds, track temporal module rewiring, and instantly validate abstract topological shifts against raw transcriptomic distributions.

Further, Tangerine’s architecture is uniquely positioned to benefit from the increasing scale of single-cell experiments. As dataset resolutions expand to hundreds of thousands of cells across denser temporal windows, the underlying metacell aggregation becomes increasingly robust. Higher cell counts natively enhance the statistical power of the aggregation step, yielding an aggregated, lower-sparsity input matrix that facilitates responsive, time-specific regularized regression. Ultimately, Tangerine accelerates the transition from computational prediction to experimental design, ensuring that dynamic regulatory inferences are mathematically sound, transparently visualized, and biologically actionable.

## Conflicts of interest

The authors declare that they have no competing interests.

## Funding

This work is supported in part by funds from the UK Research and Innovation Future Leader Fellowship (Grant/Award Number: MR/T022620/1).

## Data availability

The data underlying this article are available in CZ CELLxGENE Discover platform at https://datasets.cellxgene.cziscience.com/ae560045-adac-4c39-810a-21cec5686599.h5ad. Alternatively, the raw and processed forms of the data can be accessed from the NCBI Gene Expression Omnibus (GEO) using accession numbers GSE186069 and GSE228590.

## Author contributions statement

G.S. conceptualised the project, T.N. built the software and performed data curation. T.N. and G.S. investigated and validated the results, wrote and reviewed the manuscript.

## Acknowledgments

The authors thank Tom Owen-Hughes and Nicola Wiechens for helpful discussions. The authors thank the anonymous reviewers for their valuable suggestions.

## Notes

https://github.com/ntanmayee/tangerine

### Competing Interest Statement

The authors have declared no competing interest.

